# Islet-on-a-chip device reveals first phase glucose-stimulated respiration is substrate limited by glycolysis independent of Ca^2+^ activity

**DOI:** 10.1101/2022.03.02.482671

**Authors:** Romario Regeenes, Yufeng Wang, Anthony Piro, Aaron Au, Christopher M. Yip, Michael B. Wheeler, Jonathan V. Rocheleau

## Abstract

Pancreatic islets respond metabolically to glucose by closing K^ATP^ channels resulting in Ca^2+^-influx and insulin secretion. Previous work has revealed the importance of glycolytic flux in triggering insulin secretion. However, it is unclear whether the triggered (‘first phase’) secretion is further amplified by Ca^2+^-stimulation of mitochondrial NADH production and/or oxidative phosphorylation (OxPhos). Although commercially available tools have been developed to explore islet metabolism, these methods often overlook islet variability and have poor spatiotemporal resolution. To tease apart first phase glucose-stimulated respiration, we designed an islet-on-a-chip microfluidic device to simultaneously measure O^2^-consumption rate (OCR) and Ca^2+^-activity of individual islets with high temporal resolution. We used finite element analysis to optimize placement of sensor in optically clear microwells on a thin glass coverslip. The microfluidic channels were subsequently fabricated using O_2_-impermeable plastic to limit outside-in diffusion and push islets against the microsensor. We validated our device using living mouse islets and well-established modulators of respiration. By inhibiting glycolysis and mitochondrial pyruvate transport, we show that islet OxPhos is limited by NADH-substrate rather than ADP in low and high glucose. We subsequently imaged glucose-stimulated OCR and Ca^2+^-influx simultaneously to reveal a biphasic respiratory response that is determined by glycolytic flux through pyruvate kinase (PKM2) and independent of Ca^2+^. These data demonstrate the unique utility of our modular and optically clear O_2_-sensor to simultaneously measure glucose-stimulated OCR and Ca^2+^ activity of multiple individual islets.

## 1. Introduction

Glucose-stimulated insulin secretion (GSIS) occurs from pancreatic islets in a large first phase burst within 5 min of stimulation followed by smaller second phase oscillations (Henquin et al., 2006; Nilsson et al., 1996). GSIS is triggered from islet beta-cells by a rise in ATP/ADP ratio that closes K_ATP_ channels resulting in membrane depolarization and Ca^2+^-influx (termed ‘K_ATP_-dependent secretion’) (Bertram et al., 2006; Wang et al., 2021). Early work revealed a central role for glycolysis in triggering insulin secretion (Dukes et al., 1994; Eto et al., 1999). More recent work showed that inhibition of pyruvate kinase-M2 (PKM2) (i.e., the last step of glycolysis) abolishes insulin secretion further emphasizing the role of glycolysis in triggering insulin secretion (Nakatsu et al., 2015). Insulin secretion is further amplified by glucose-metabolism independent of K_ATP_ channel activity (termed ‘K_ATP_-independent amplification’) effectively doubling insulin secretion (Henquin, 2009; Mourad et al., 2010). The amplification of second phase insulin secretion has been classically linked to mitochondrial anaplerosis and associated efflux of metabolic intermediates (Campbell and Newgard, 2021) as well as Ca^2+^-induced mitochondrial metabolism and respiration (Quan et al., 2015; Tarasov et al., 2012). Whether these same mechanisms also amplify first phase insulin secretion is less defined.

It is well recognized that islet mitochondrial metabolism and OxPhos are key determining factors of GSIS that are significantly impaired in type 2 diabetes (T2D) (Antinozzi et al., 2002; Haythorne et al., 2019). O_2_-consumption rate (OCR) is a particularly informative parameter of mitochondrial function as it reflects both electron transport chain (ETC) activity and NADH-substrate supply (Will et al., 2007). Thus, measuring first phase OCR could help reveal the role of mitochondrial metabolism in amplifying first phase insulin secretion. The lack of studies examining amplification of first phase insulin secretion is likely due to technological limitations of simultaneously measuring OCR (related to OxPhos) and Ca^2+^ (insulin secretion triggering and mitochondrial activation). Specifically, such measurements would need sufficient temporal resolution to discern the first (< 5 min) and second phases of insulin secretion. In addition, these measurements should ideally be performed simultaneously on single islets to avoid blurring of the temporal responses due to islet-to-islet variability (Taddeo et al., 2018).

Various methods are commercially available to measure islet respiration. However, these methods often do not provide optical access to simultaneously measure Ca^2+^ activity, and when they do, they lack sufficient spatial and temporal resolution to measure individual islet responses (Nászai et al., 2019; Taddeo et al., 2018). To address these limitations, we explore placing optical O_2_ sensor inside microfluidic devices previously designed for quantitative live cell imaging of pancreatic islets (Sankar et al., 2011; Silva et al., 2016, 2013). These devices hold islets against a thin glass coverslip, which provides an ideal location to place optical sensors to measure OCR while simultaneously imaging Ca^2+^-influx. We subsequently use this device to demonstrate that first phase respiration is NADH-substrate limited by glycolytic flux and independent of Ca^2+^ activity.

## 2. Materials and Methods

### 2.1 Pancreatic islet isolation and culture

Animal procedures were approved by the Animal Care Committee of the University Health Network, Toronto, Ontario, Canada in accordance with the policies and guidelines of the Canadian Council on Animal Care (Animal Use Protocol #1531). Pancreatic islets were isolated from 12-14 weeks old C57BL/6 male mice as previously described (Sankar et al., 2011; Silva et al., 2013) and cultured overnight in a humidified incubator (37°C, 5% CO_2_).

### 2.2 Microfluidic device fabrication

The custom microfluidic device comprised a ‘top piece’ and glass coverslip with O_2_-sensor microwells. The top piece with 80 µm tall channels and 60 µm tall dam-wall was fabricated using 3 mm thick polymethylmethacrylate (PMMA) and a Nomad 883 Pro mill. A 1/16” endmill was used to mill the device followed by a 1/8” µm endmill for the inlet and outlet ports. The top piece was washed in 70% ethanol and left to dry before attachment to the microwell coverslip. Microwells were fabricated onto no. 1.5 glass coverslips using SU-8 photoresist (Micro-Chem) exposed through a transparent plastic film. The O_2_-sensor was prepared by mixing polydimethylsiloxane (PDMS) (10:1 ratio) and Platinum(II)-5,10,15,20-tetrakis-(2,3,4,5,6-pentafluorphenyl)-porphyrin (PtTFPP) (0.1 mg/1 mL in toluene) at a volume ratio of 10:1 (Thomas et al., 2009). 100 µL of this mixture was left to dry (30 min) prior to squeegeeing into the microwells. The slide was subsequently left to dry in the dark (24 h at room temperature). The top and glass coverslip were carefully assembled using epoxy-based glue and clamped together (1 h).

### 2.3 Microfluidic fluid flow modelling/oxygen sensor

The device was constructed in two-dimensions using AutoCAD 2020 prior to being extruded to three-dimensions in COMSOL Multiphysics 5.4 (COMSOL). The fluid flow was studied using the laminar flow model and dissolved O_2_ was studied using the transport of diluted species model. Pancreatic islets were modelled as 120 µm radii cylinders with the circumferential edge filleted and modeled to consume O_2_ under a maximum rate of 0.034 mol·s^-1^·m^-3^ (Buchwald, 2009). Islets were modelled as a flattened sphere/cylinder due to how the islets are pushed up against the sensor and ceiling of the device. To determine O_2_-concentration underneath the islet, a *cut line surface normal* was used with the same coordinates for all the simulations. Inlet pressure for all the simulations was set to 0 kPa while the exit flow rate was set to 200 µL·h^-1^, unless otherwise stated. The simulated models had meshes with 50735-174199 domain elements, 11221-14864 boundary elements, and 818-11184 edge elements. All diffusion coefficients were used at 37°C for our numerical simulations.

### 2.4 Microscope setup

Luminescence imaging was done using the 40×/0.75 NA air objective of an inverted widefield RAMM fluorescence microscope (ASI) equipped with 3 LEDs (460, 535, and 560 nm), stage top incubator (Okolab) and Iris 15 sCMOS camera (Photometrics). Images were binned resulting in 1264 × 740 pixels at 0.425 µm/pixel. The microscope was controlled using MicroManager (Edelstein et al., 2014). PtTFPP was imaged using two 500 ms bursts of the 535 nm LED and a 590-660 nm band-pass emission filter. Cal-520 fluorescence was excited using 300 ms of 465 nm LED exposure attenuated to 32% and collected through a 495-525 nm band-pass emission filter.

### 2.5 Sensor calibration

The PMMA device with O_2_-sensing microwells was subjected to flowing solutions of varying dissolved O_2_ concentrations. To measure the Stern-Volmer relationship of the PtTFPP/PDMS sensors, different concentrations of O_2_ were set in the device by flowing solution from a 1L volume of ddH_2_O enclosed in a glass chamber that was progressively treated with increasing amounts of sodium sulfite (10 % at 5000 µL·h^-1^). The decreasing concentration of dissolved O_2_ in the chamber was measured every two minutes relative to the sensor response using a miniDOT USB oxygen logger (PME). The mean luminescence intensity of the on-chip PtTFPP/PDMS sensor *versus* O_2_ concentration was modelled as a linear regression using the Stern-Volmer equation:

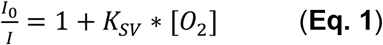

where *I*_*0*_ and *I* are the luminescence intensity in the absence and presence of molecular oxygen, *[O*_*2*_] and *K*_*SV*_ is the Stern-Volmer constant.

### 2.6 Glucose stimulated oxygen imaging

Islets were pre-incubated 1 h in low glucose (2 mM) imaging buffer (125 mM NaCl, 5.7 mM KCl, 0.42 mM CaCl_2_, 0.38 mM MgCl_2_, 10 mM HEPES, and 0.1% BSA; pH 7.4) at 37°C. Islets (2 to 5 at a time) were subsequently loaded into a stage-mounted microfluidic device (37°C) with flow (200 µl·h^-1^) due to pull from a syringe pump. The islets were exposed to various treatments as indicated by exchange of the media in an on-chip well prior to luminescence imaging.

### 2.7 Simultaneous OCR- and Ca^2+^-imaging

Islets were pre-incubated in imaging buffer with 4 μM Cal-520 (Life Technologies) (1 h, 37°C, 2 mM glucose, and 1 mM N-acetylcysteine (NAC)) (Paz-Miguel et al., 1999). The islets were subsequently loaded into a stage-mounted microfluidic device and pre-incubated 10 min in flowing imaging buffer (200 µl·h^-1^, 37°C, 2 mM glucose, 1 mM NAC) prior to imaging. A custom macro was created to collect PtTFPP and Cal-520 images using 535 and 460 nm excitation every 10 s, and at two different z-planes to account for the difference in imaging plane between the microwells and Cal-520 loaded islets. The islets were then variably treated as indicated by replacing media in the on-chip well while imaging.

### 2.8 Image analysis

All images were analyzed using ImageJ. Regions of interest (ROI) were manually drawn around each microwell underneath the islet. After selection, the mean intensity for each microwell was recorded after background correction. The O_2_-concentration in each microwell was subsequently determined using the K_SV_ of the calibrated microsensors by rearranging Eq. 1. OCR of individual islets was calculated using the following equation:

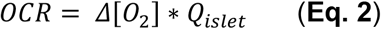

where *Δ[O*_*2*_] is the average change in O_2_-concentration of the microwells underneath the islet and *Q*_*islet*_ is volumetric flow rate across the islet, approximating the islet as a flattened sphere. The dissolved O_2_-concentration was calculated to be 207.3 µM at 37 °C with no islet present in the device. Ca^2+^ images were analyzed by normalizing the fluorescence intensity of randomly selected ROIs by the mean intensity during 2 mM glucose treatment.

## 3. Results

### 3.1 Design and optimization of an islet-on-a-chip device with modular O_2_-sensors

Our previous islet-on-chip devices offered ready placement of optical O_2_-sensors (Sankar et al., 2011; Silva et al., 2016, 2013). To explore placing sensor underneath the islets, we modelled islets in a dam-wall microfluidic device (**Fig. 1**). We chose an islet-on-a-chip device that immobilizes islets against the glass coverslip in flowing media using a drop in channel height (80 to 20 µm) / dam wall (**Fig. 1A**). The drawing (not to scale) shows a device fabricated out of O_2_-impermeable PMMA (grey) with PtTFPP sensor embedded in O_2_-permeable PDMS (yellow) that coats the glass coverslip. We normally fabricate microfluidic channels out of PDMS but switched to PMMA to avoid swamping/masking islet O_2_-consumption. We modelled both an empty device (left) and islet-filled-device (right) using finite element analysis to determine O_2_ concentration in the sensor layer (**Fig. 1B**) including along the centerline plotted with flow from left to right (**Fig. 1C**). The two concentric circles (**Fig. 1B, right**) represent the outer diameter of the islet, and the smaller footprint of the islet pressed against the sensor. These data show limited response underneath the islet (right image, red and blue circles) consistent with fast O_2_-diffusion along the PDMS substrate dampening the response to islet respiration. To limit this diffusion, we subsequently modelled the sensor in microwells (**Fig. 1D-H**). The drawing (not to scale) depicts microwells formed using O_2_-impermeable SU-8 (black), which limits diffusion between the microwells (yellow) (**Fig. 1D**). To determine the impact of microwell size, we modelled the OCR of islets placed against microwells of varying widths (10, 50 and 100 µm) (**Fig. 1E-F**). These data show a progressively larger response with smaller microwell width. The microwells are more sensitive once they are fully covered by the islet footprint. Microwells that are partially uncovered (due to large size or being on the edge of the footprint) are susceptible to fast O_2_ diffusion around the tissue and through the PDMS. We also simulated 10 µm wide microwells with varying depths (10, 30 and 60 µm) (**Fig. 1G-H**). These data show no difference in the response based on microwell depth. To further evaluate the impact of the SU-8 layer on imaging quality, we imaged fluorescent beads on different thicknesses of SU-8 **(Fig. S1)**. These data show a progressive loss of fluorescence intensity with increasing SU-8 thickness. From these data, we chose to fabricate microwells that are 10 µm wide with a 20 µm pitch and 10 µm tall. Furthermore, we modeled the effects of flow rate, islet proximity to sensor surface, and variable oxygen consumption rates (**Fig. S2**). Overall, these data show that flow rates do not significantly impact sensing (**Fig. S2A**), islets need to be up against the glass coverslip to maximize the response (**Fig. S2B and C**), and our microsensors could measure changes in consumption from low to high glucose (**Fig. S2D**).

**Figure 1.**
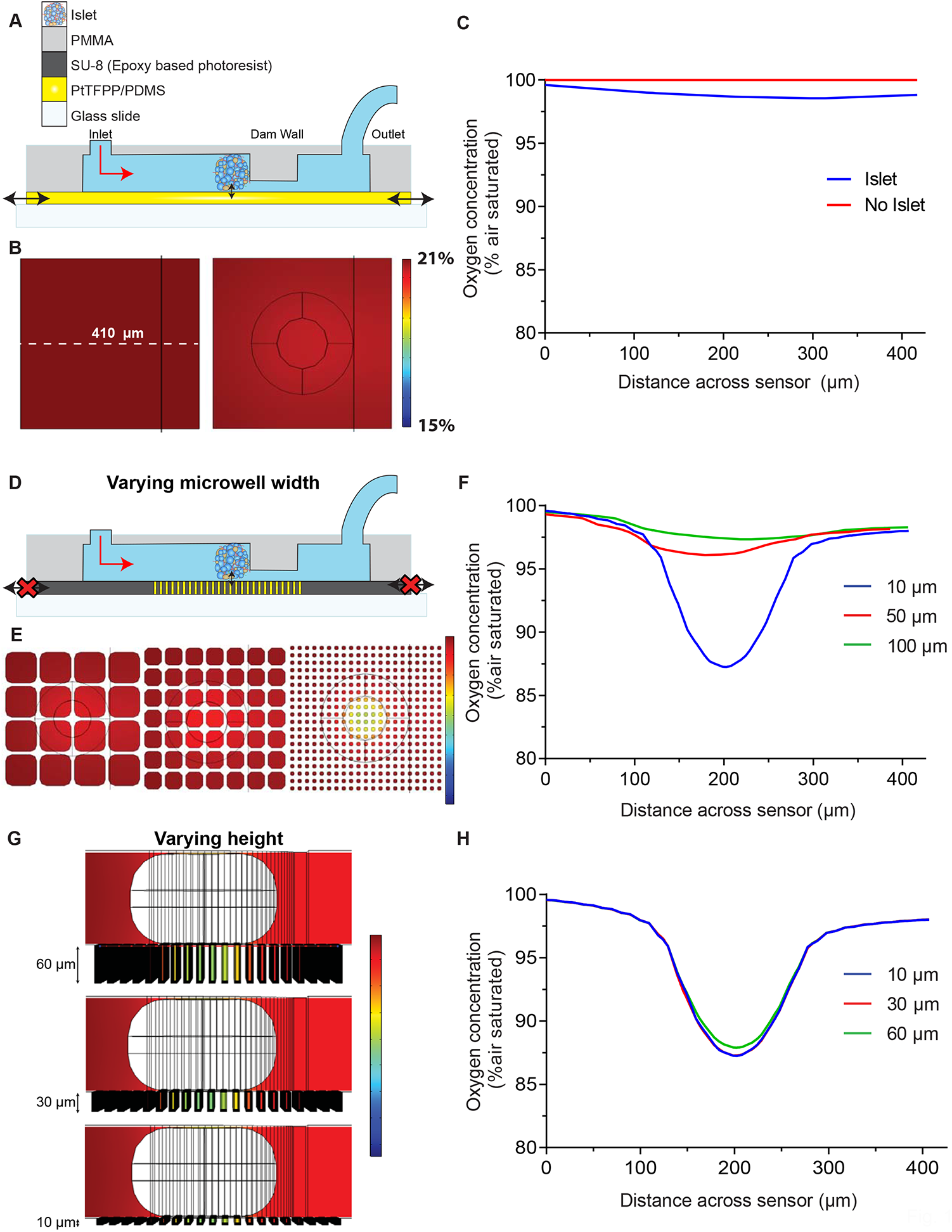
COMSOL simulations reveal greater sensitivity when patterning O_2_ sensor in microwells. A) A schematic of various entry points for external O_2_ into the microfluidic device. The planar sensor made from PDMS allows O_2_ from the environment to enter and leave lowering sensor sensitivity directly beneath the islet. Red arrow indicates direction of fluid and O_2_ flow. Black arrows indicate regions where O_2_ can enter and leave. B) *Top-down view*: O_2_ transport in the device in the absence *(left)* and presence (*right)* of an O_2_ consuming islet with the PtTFPP/PDMS in a uniform plane underneath. White dotted line represents distance across sensor where 0 denotes the far left (µm). C) Simulated O_2_ concentrations across the PtTFPP/PDMS in the presence of an islet reveal a limited response across the planar sensor. D) A schematic with O_2_ entry points sealed by patterning SU-8. This arrangement digitizes the sensor readout and prevents O_2_ from travelling between wells.E) *Top-down view*: O_2_ concentration distributions in 10 (*right*), 50 (*middle*) and 100 (*left*) µm wells with an islet above. F) Simulated O_2_ concentrations in 10, 50 and 100 µm wells underneath the islet reveal a dramatic increase in sensitivity with 10 µm wells. G) *Side view*: O_2_ concentration distributions in 10 (*bottom*), 30 (*middle*) and 60 (*top*) µm height wells reveal no changes in sensor response. H) Simulated O_2_ concentrations in 10, 30 and 60 µm height wells reveal no change in OCR sensitivity *(right)*. Pancreatic islets were modelled as 120 µm radii cylinders with the circumferential edge filleted.

### 3.2 Imaging OCR in individual pancreatic islets

To measure OCR of living islets, we fabricated a dam wall microfluidic device to trap and image islets on top of the optical microsensor (**Fig. 2A**). This device included an on-chip well for islet loading through an inlet and fast exchange of media, a dam wall to trap the islets, and an outlet tube connected to a syringe pump to control flow rates. This design traps islets of varying shapes and sizes either in the channel prior to the dam wall (>80 µm diameter) or under the dam-wall (∼20 µm diameter). To estimate the impact of flow on islet function, we modelled shear stress on the surface of an islet next to the dam-wall of the device at a flow rate of 200 µl·h^-1^ (**Fig. S3**). These data indicate that most of the shear stress is below 5 mPa similar to previous devices that maintained glucose-stimulated Ca^2+^ activity even after 48 h of flow (Silva et al., 2013). O_2_ sensors are generally calibrated prior to use so that luminescence intensities can be related back to O_2_ concentrations using the Stern-Volmer relationship (Vanderkooi et al., 1987). To determine our sensor dynamic range, we flowed double distilled water (ddH_2_O) and an O_2_-scavenger (10% sodium sulfite) to obtain luminescence intensities for 100% air saturated and 0% O_2_, respectively (**Fig. 2B**). These images show an ∼25-fold change in microsensor intensity in the presence of an O_2_-scavenger. To characterize the on-chip O_2_ sensor, we determined the Stern-Volmer relationship, response time (t_95_) and sensor repeatability **(Fig. S4**). Overall, these data indicate that our sensor responds linearly with a limit of detection of ∼51 nmol·L^-1^, response time of <15 s, and 5.8 % relative standard deviation between devices. We subsequently loaded living mouse pancreatic islets in our device to measure the dynamics of OCR. Once loaded, islets sitting on the microsensor could be imaged by an inverted widefield epifluorescence microscope (**Fig. 2C**). Islet bioenergetics are commonly measured in batch (∼70 islets) using commercially available devices (e.g., Seahorse XF analyzer) and a series of mitochondrial respiration treatments including: (i) 2 mM glucose to measure basal OCR, (ii) 20 mM glucose to measure glucose-stimulated OCR, (iii) oligomycin to measure ATP-linked respiration, (iv) FCCP to measure maximal respiratory capacity, and (v) rotenone + antimycin to define non-mitochondrial respiration. To validate our device, we used these same treatments on individual islets (**Fig. 2C**). The microsensors underneath the islet showed clear changes in luminescence intensities reflecting their respective measurements of O_2_. These data show that the O_2_ sensor can measure pharmacologically induced changes in OCR in individual pancreatic islets. However, unlike commercial assays, we are not limited to a narrow size of islets (Taddeo et al., 2018) (**Fig. 2D**). To determine the effect of variable islet size on the OCR, we plotted the basal OCR relative to islet volume estimated from the widefield image (**Fig. 2E**). These data show a linear correlation between islet volume and OCR (R^2^ = 0.92, p<0.0001). We therefore normalized the individual islet responses to the islet volumes prior to plotting the responses to the respiratory inhibitors (**Fig. 2F**). These data show a 2-fold increase from basal to glucose-stimulated OCR and maximal respiration consistent with responses from batch analysis using commercially available instruments (Taddeo et al., 2018) (**Fig. 2G**). These data also show the uncoupled state (i.e., proton leak) is approximately 68% of basal OCR consistent with previous studies (Wikstrom et al., 2012). Overall, these data validate that our device can measure OCR of individual pancreatic islets.

**Figure 2.**
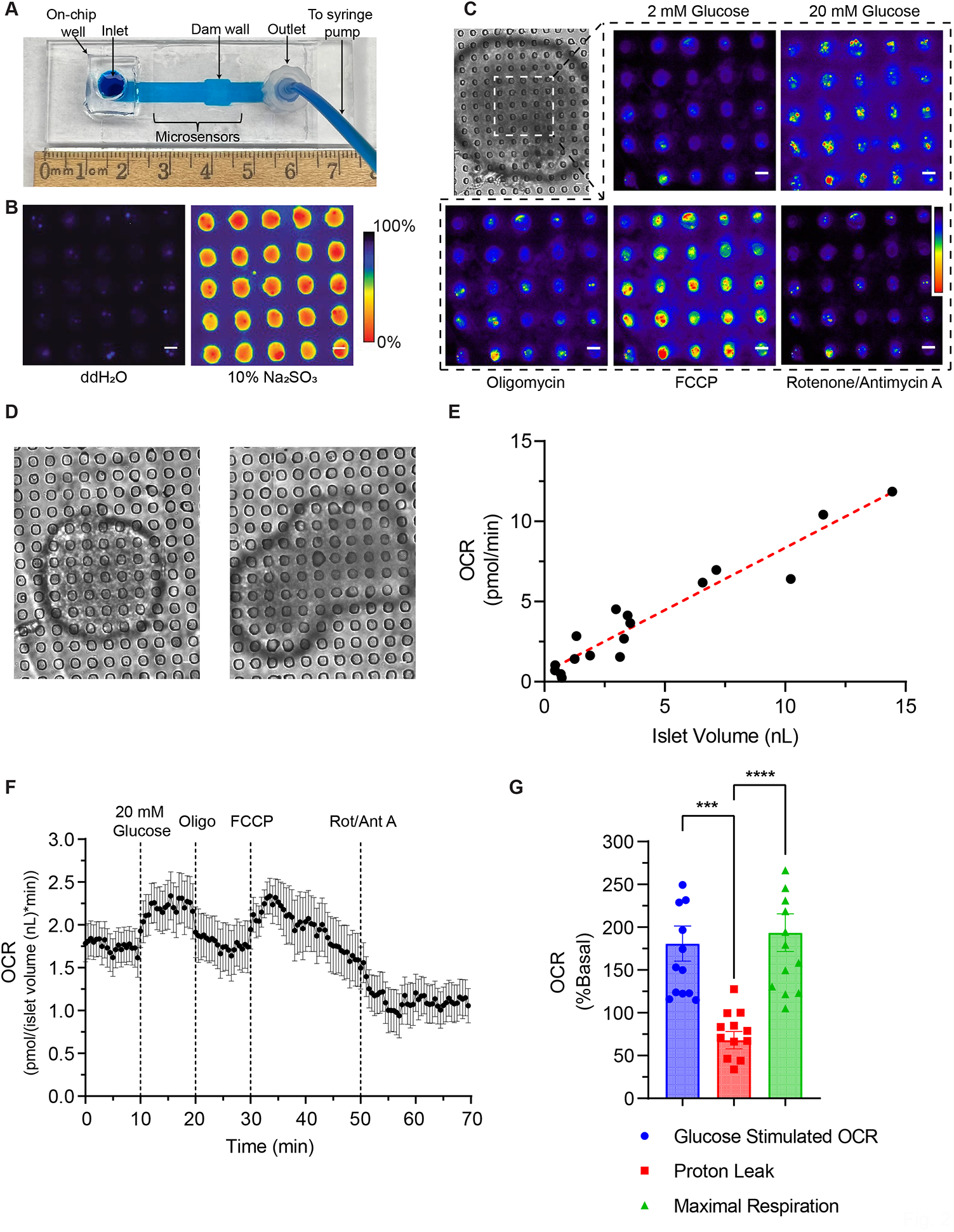
An on-chip O_2_ sensor to measure OCR in individual pancreatic islets. A) Photograph of final microfluidic device filled with blue dye for easy visualization. B) O_2_ sensitive microwells in the presence of ddH_2_O (*left*) and 10% sodium sulfite (*right*). C) Representative widefield phosphorescence images of O_2_ sensitive microwells underneath an islet while flowing the indicated respiratory inhibitors. D) Representative widefield white light images of different sized islets trapped in the device. E) Basal OCR of individual mouse islets (*black dot*) relative to islet volume. N = 17 islets. F) Mean OCR of mouse islets exposed to 20 mM glucose, 5 µM oligomycin, 2 µM FCCP and 5 µM rotenone/antimycin A. G) Islet bioenergetic measurements extracted from the respiratory inhibitor treatments. Glucose stimulated OCR, proton leak and maximal respiration relative to basal OCR. N = 14 islets. Data in figure are means ± SEM. **** indicates p ≤ 0.0001 and *** indicates p ≤ 0.001 by one-way ANOVA. Scale bars represent 10 µm.

### 3.3 Temporal imaging of the glucose-stimulated OCR and Ca^2+^ responses

Glucose-stimulated OCR has previously been shown to rise prior to Ca^2+^-influx suggesting that Ca^2+^ does not play an immediate role (Jung et al., 2000). To determine the temporal contribution of Ca^2+^ to glucose-stimulated metabolism, we loaded mouse pancreatic islets with Cal-520 to simultaneously image OCR and Ca^2+^ activity in response to glucose (**Fig. 3A and B**). The Ca^2+^ traces upon glucose stimulation show a characteristic ‘phase 0’ drop upon glucose stimulation due to activation of Ca^2+^-ATPase ER pump efflux and ‘phase 1’ rise ∼3 min after stimulation due to voltage-gated Ca^2+^ channel influx (**Fig. 3B**). Notably, the glucose-stimulated OCR traces were biphasic showing a simultaneous rise with the drop in Phase 0 Ca^2+^ and a second rise near or before phase 1 Ca^2+^-influx. To determine the magnitude of the OCR responses, we calculated the normalized OCR area under the curve (AUC) for phase 0 and phase 1 as defined by the individual Ca^2+^ traces (**Fig. 3C and D**). In other words, we used the changes in Ca^2+^-activity to define the 2mM glucose, phase 0, and phase 1 regions for each OCR trace (**Fig. 3C**). Using this criterion, the OCR increased significantly and sequentially from basal to phase 0 and to phase 1 (**Fig. 3D**). To further explore the biphasic OCR response, we focused on the relative changes in OCR within the first two minutes of phase 0 and phase 1 by setting “Time 0” for each curve based on the inflection points of the Ca^2+^ traces (**Fig. 3E and F**). Due to variability in the timing of the responses, this analysis could only be done due to our ability to measure traces from individual islets (**Fig. S5**). The first rise in OCR occurs simultaneously with the phase 0 drop in Ca^2+^ plateauing within 1 min and prior to phase 1 Ca^2+^-influx (**Fig. 3E**). These data are consistent with an increase in OxPhos due to the metabolic flux (NADH supply) that is independent of mitochondrial stimulation by Ca^2+^. The initial plateau was followed by a second rise in OCR that preceded Ca^2+^-influx (**Fig. 3F**). Further analysis of the OCR curve using linear regression of the eight time points starting from time = 0 min fit to an x-intercept of -27.4 ± 10.9 s. In other words, the second OCR response preceded Ca^2+^-influx by ∼28 s. To further confirm the biphasic OCR response to glucose, we measured the slope of the OCR response for each trace before and after the -30 s inflection point **(Fig. 3G)**. These data show significantly different slopes once again consistent with a further inflection in OxPhos. Overall, these data illustrate that islet respiratory response to glucose is biphasic with the step in the response occurring before and independently of Ca^2+^-influx.

**Figure 3.**
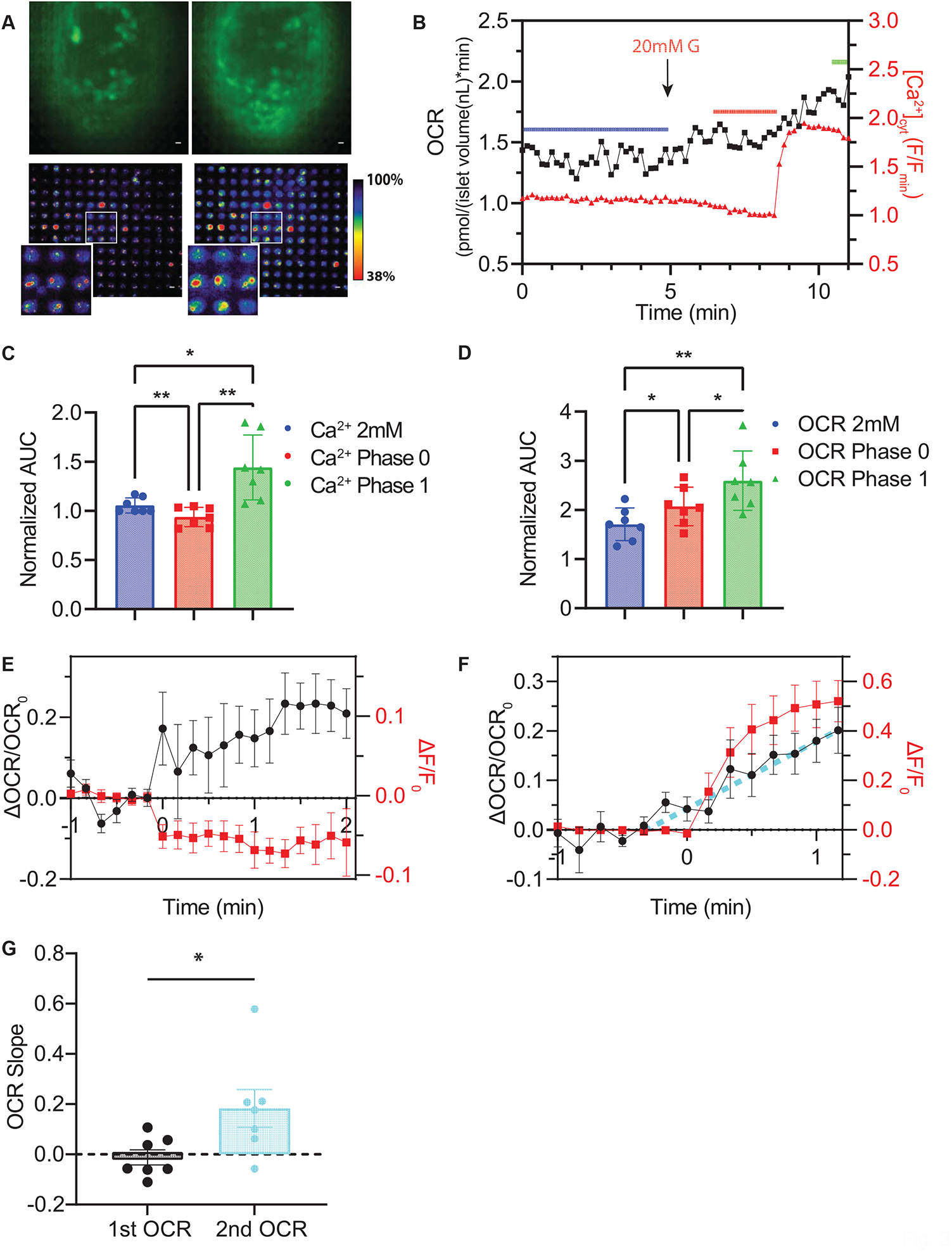
Dual-colour imaging of OCR and Ca^2+^ in individual islets reveals a two-step increase in OCR response. A) Representative images of a Cal-520 stained mouse islet and O_2_ sensitive microwells underneath the islet after stimulating with 2- (*left*) and 20-mM glucose (*right*) show a visible increase in luminescence in both sensors. Scale bars represent 10 µm. B) Representative OCR and Ca^2+^ traces of an individual mouse islet show OCR rise precedes Ca^2+^ influx. The black arrow indicates treatment change from 2-to 20-mM glucose. The Ca^2+^ response was normalized to the Cal-520 fluorescence intensity during low glucose treatment. C-D) Averaged area under the curve (AUC) of Ca^2+^ (C) and OCR (D) traces during low glucose treatment, phase 0 and 1 Ca^2+^ responses. Analyzed areas are indicated by the blue, red and green lines in B). E-F) Average normalized changes in OCR determined by setting “time = 0 min” based on the start of 20 mM glucose stimulation (E) and the beginning of phase 1 Ca^2+^ response (F). Changes in OCR were normalized to the six time points prior to t = 0 min. Cyan line in F) indicates the fitting for the start of the second OCR rise. G) Averaged slopes of the two-step OCR responses to high glucose stimulation. Data in figure are means ± SEM. ** indicates p ≤ 0.01 and * indicates p ≤ 0.05 by one-way ANOVA. N = 7 islets.

### 3.4 PKM2 is responsible for the biphasic rise in OCR

The rate of OxPhos depends on the metabolic supply of NADH (‘substrate-limited’) or when NADH is in excess by the availability of ADP (‘ADP-limited’) (**Fig. 4A**). We postulated that the biphasic glucose-stimulated OCR response reflects islets in a substrate-limited state with transitions in OCR reflecting greater production of NADH by glycolysis and the TCA cycle. To first confirm islet OCR is substrate-limited at low glucose, we measured the OCR response of islets in low glucose to inhibition of glycolytic flux using 2-deoxyglucose (2DG) and in response to blocking entry of pyruvate into mitochondria using UK5099 (**Fig. 4B**). These data show 2DG did not induce a change in OCR consistent with low glycolytic flux when glucose is below the K_m_ for glucokinase. However, UK5099 induced a small but significant decrease in OCR indicating that basal respiration is dependent on pyruvate entry. These data highlight the importance of pyruvate entry into the TCA cycle in stimulating OxPhos (3 NADH/pyruvate) and potentially reflect amino acid oxidation (e.g., alanine, serine) at low glucose. To confirm that glucose-stimulated OCR is due to glycolytic flux and substrate limited, we measured the OCR response of glucose-stimulated islets to 2DG (**Fig. 4C and D**). These data show that glucose-stimulated OCR dropped to basal (low glucose) levels by inhibition of glycolytic flux again suggesting islet respiration ultimately depends on substrate supply. Due to the varied 2DG and UK5099 responses in low glucose, we speculated that the biphasic nature of the glucose-stimulated OCR reflects downstream regulation of PKM2, which is allosterically regulated by the glycolytic intermediate fructose 1,6-bisphosphate (FBP) (Ashizawa et al., 1991). To investigate the role of PKM2 in setting basal OxPhos, we measured the OCR of islets treated for 1 h with PKM2 activator TEPP-46 (**Fig. 4E**). These data show a small but significant increase in basal OCR induced by PKM2 activation again consistent with OCR being substrate limited (in this case by the supply of pyruvate from glycolysis). To further investigate the role of PKM2 activity in the glucose-stimulated response, we measured the glucose-stimulated OCR response of TEPP-46 treated islets (**Fig. 4F**). Glucose induced a sharp decrease in OCR in the presence of TEPP-46. This rapid reduction in OCR is consistent with the rapid entry of pyruvate due to fully active PKM2 leading to lower NADH induced by lactate dehydrogenase activity, conversion to oxaloacetate/malate through pyruvate carboxylase activity, and/or depolarization of mitochondrial membrane potential. Notably, the OCR subsequently increased out to 15 min of stimulation with no sign of a biphasic response. In other words, the biphasic glucose-stimulated response observed in untreated islets is replaced by a single faster rise. To determine the role of Ca^2+^-influx in this response, we simultaneously imaged the glucose-stimulated OCR and Ca^2+^ responses of TEPP-46 treated islets (**Fig. 4G**). These data again show a sharp drop in OCR that coincides with the beginning of phase 0 Ca^2+^-efflux followed by a linear rise in OCR that noticeably precedes phase 1 Ca^2+^-influx. Normalizing the curves from several islets relative to glucose arrival showed the linear increase in OCR precedes Ca^2+^-influx by ∼3.2 min (**Fig. 4H**). Collectively, these results confirm islet OCR is substrate-limited by glycolytic flux and suggest that PKM2 plays a critical role in regulating early glucose-stimulated OCR.

**Figure 4.**
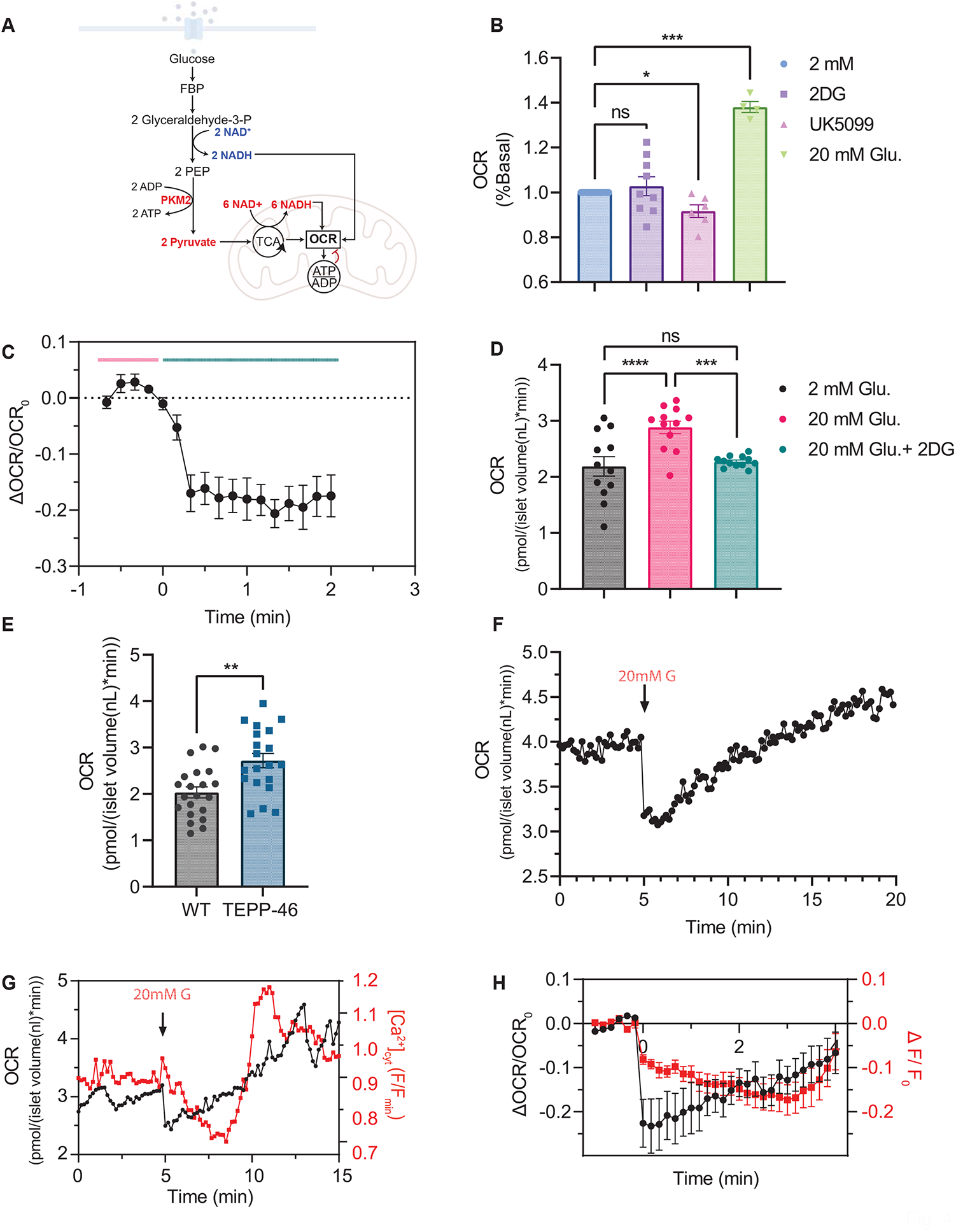
Glycolytic flux and pyruvate entry into mitochondria control OCR. A) Simplified metabolic model depicting the impact of glycolysis derived NADH and PKM2 activity on OCR. NADH from glycolysis is shuttled into the mitochondria for charging the electron transport chain. PKM2 controls pyruvate flux into the mitochondria for tricarboxylic acid (TCA cycle) metabolism, which also generates NADH to further fuel the electron transport chain. Substrate (NADH) supply establishes a proton gradient stimulating OCR due to OxPhos/ATP production. In excess substrate supply, OCR will be inhibited by a rise in the ATP/ADP ratio. B) Normalized basal OCR rates in the presence of 2DG, a glycolysis inhibitor, and UK5099, a mitochondrial pyruvate carrier inhibitor and 20 mM glucose. C) Normalized changes in OCR of islets in 20 mM glucose in response to 20 mM + 2DG reveals substrate supply is needed to stimulate OCR. D) Averaged OCR from 3-minute intervals during steady state for each treatment. E) Preincubating islets with PKM2 activator (4 μM TEPP-46) shows a significant increase in basal OCR levels consistent with greater pyruvate flux.F) A representative trace of a single islet preincubated in 4 μM TEPP-46 responding to 20 mM glucose. G) A representative OCR and Ca^2+^ trace of an individual islet TEPP-46 treated shows OCR precedes Ca^2+^-influx. H) Individual islet traces were normalized to the onset of 20 mM glucose stimulation to reveal a rapid drop followed by linear increase in OCR prior to Ca^2+^-influx. Data in figure are means ± SEM. **** indicates p ≤ 0.0001, *** indicates p ≤ 0.001, ** indicates p ≤ 0.01, and * indicates p ≤ 0.05 by one-way ANOVA. N = 4-21 islets.

## 4. Discussion

Our results demonstrate the unique capability of our islet-on-a-chip microsensor to measure OCR and Ca^2+^-activity of multiple individual islets (3-5 / run) with high temporal (<15 s) resolution. This was achieved by pushing islets against an optically clear digitized microsensor. In comparison, Clark electrodes are unable to measure OCR of individual islets since they suffer from gradual consumption of O_2_ (Wolfbeis, 2015). The Seahorse analyzer provides insufficient temporal resolution of single islet responses (>5 min) and cannot simultaneously measure Ca^2+^ activity (Taddeo et al., 2018). The Oroboros O2k analyzer is capable of measuring OCR and Ca^2+^ simultaneously but does so in a bulk measurement that blurs the highly variable responses of individual islets (Morota et al., 2013). Finally, microelectrode sensors allow simultaneous OCR- and Ca^2+^-measurements; however, they require islets to be cultured onto a dish (i.e., not freshly isolated), use highly specialized equipment, and in comparison, are low in islet throughput (Jung et al., 2000).

Our microsensor is sensitive to nanomolar changes in O_2_, shows fast response times (<15 s) and is easy to calibrate. To achieve these characteristics, we embedded optically responsive sensor (PtTFPP) within an O_2_ permeable substrate (PDMS) based on previous work showing this combination had high photostability, large Stokes shift and dynamic response (Lee and Okura, 1997; Thomas et al., 2009). Conventional optical O_2_ sensors for microfluidic devices are generally planar and open to the environment. Our initial modelling suggested that O_2_ diffusion inside the planar sensor masks OCR responses of individual islets. Thus, we placed the PtTFPP/PDMS sensor into micron-sized wells surrounded by O_2_ impermeable epoxy. We improved the sensitivity by reducing the microwell width to less than the footprint of islets (∼50 µm in diameter) and showed the importance of using microchannels to push the tissue against the sensor. We subsequently used PMMA to fabricate the channels (i.e., the top of the device) instead of more conventionally used and O_2_-permeable PDMS. Fabricating the device using air saturated PDMS abolished the sensor response during early trials consistent with O_2_ diffusion from the air saturated polymer swamping the sensor response.

Static OCR responses using our device were consistent with previous work once we normalized our data against individual islet volume (Taddeo et al., 2018). We showed that islet OCR is highly uncoupled from ATP production using oligomycin consistent with other bulk islet studies (**Fig. 2G**) (Taddeo et al., 2018; Wikstrom et al., 2012). A unique feature of our microsensors is that they are fabricated using a thin glass coverslip and are optically clear, both of which allow simultaneous imaging of other responses. We used our microsensors to measure the OCR responses of individual islets while simultaneously imaging Ca^2+^-activity. Our microsensor shows high temporal resolution due to digitization of the sensor. In contrast, many labs use commercial instruments that are limited to a bulk measurement every 2-5 minutes, thereby masking islet dynamics, and prohibiting multiparametric imaging. These approaches are unable to acquire data at earlier timepoints and thus may not capture evidence for respiratory control by NADH substrate supply prior to ATP demand.

We showed OxPhos is substrate-limited throughout first phase glucose-stimulated insulin secretion (**Fig. S6**). In low glucose (i.e., below the K_m_ of glucokinase), respiration was unchanged by 2DG and diminished by inhibiting mitochondrial pyruvate transport. These data suggest OxPhos in low glucose is NADH-rather than ADP-limited and set by a slow steady-state entry of pyruvate into mitochondria. Glucose stimulation resulted in a biphasic increase in respiration with the first response plateauing within 1 min of glucose-stimulation and a second steady rise in respiration initiated ∼28 s prior to Ca^2+^-influx (**Fig. S6A**). There was no obvious change in the OCR slope during or even shortly after Ca^2+^-influx suggesting no change in NADH supply and/or OxPhos. We postulate that the initial plateau is due to glycolytic NADH (**Fig. S6A, lower left**). If islets were in an ADP-limited state, we would expect glycolysis (which produces 2 net ATP / glucose) to decrease OxPhos. Furthermore, we postulate that the plateau represents maximal flux of glycolysis through ‘inactive’ PKM2 and the subsequent linear rise in OCR is due to allosteric activation of PKM2 (**Fig. S6B, lower right**). This conclusion is based on the TEPP-46-induced rise in basal OxPhos and earlier linear rise in glucose-stimulated OCR fully separated from Ca^2+^ influx. PKM2 is heavily expressed in pancreatic islets (Martens et al., 2010) and tightly regulated by numerous posttranslational mechanisms including phosphorylation and acetylation (Prakasam et al., 2018) and by allosteric activation by FBP (Ashizawa et al., 1991). Thus, PKM2 is expected to transition from being inactive (i.e., low pyruvate flux) in low glucose to being active (i.e., high pyruvate flux) in high glucose (Ashizawa et al., 1991). The immediate glucose-stimulated dip in OCR in the presence of TEPP-46 could be due to pyruvate flux into mitochondria resulting in depolarization of mitochondrial membrane potential (De Andrade et al., 2004) and/or flux of pyruvate into mitochondrial metabolism through PC (Rocheleau et al., 2002). We postulate that reaching maximal glycolytic NADH would offset this pyruvate-stimulated dip in mitochondrial respiration. Alternatively, ATP produced by PKM2 could also decrease OxPhos as shown to regulate second phase secretion (Lewandowski et al., 2020); however, the speed of the dip is inconsistent with control by cytoplasmic ATP generation and our evidence that NADH supply determines OxPhos prior to stimulation. Consistent with a two-step glycolytic flux model, Patterson et al. previously showed a stepped rise in glucose-stimulated islet NAD(P)H response that due to the short time-lag between the steps (<20 s) suggesting the stimulus was not triggered by Ca^2+^-influx but rather stimulated by entry of pyruvate into the TCA (Patterson et al., 2000). Overall, our data are consistent with a two-step response to glucose that is due to Ca^2+^-independent activation of PKM2 resulting in pyruvate flux into mitochondrial metabolism.

## 5. Conclusions

We redesigned a dam wall microfluidic device to hold multiple pancreatic islets (3-5 / run) directly against a digitized optical O_2_-microsensor. The optical clarity and fabrication of the microwells allowed simultaneous Ca^2+^-imaging. Through the application of our islet-on-a-chip, we revealed a biphasic OCR model independent of Ca^2+^-influx and identified a role for PKM2 in controlling the biphasic behavior. While this study successfully imaged both OCR and Ca^2+^, there were some limitations. We used 10 µm thick SU-8 photoresist for rapid prototyping in the present study. This diminished the optical throughput and image quality somewhat due to SU-8 having a slightly larger refractive index than glass (∼1.6 vs. 1.5) (Ashraf et al., 2019). Future devices will be fabricated using a 3D glass printer to achieve optimal optics and thus improve dual color imaging (Burshtein et al., 2019). Overall, we anticipate this device will lead to many more applications including simultaneous imaging with other spectrally resolved sensors (Sanford and Palmer, 2017), and clinical assessment of donor islets prior to transplantation for the treatment of type 1 diabetes (Papas et al., 2015; Shapiro et al., 2006).

## CRediT authorship contribution statement

**Romario Regeenes:** Conceptualization, Methodology, Validation, Formal analysis, Investigation, Writing – Original Draft, Writing – Review & Editing, Visualization **Yufeng Wang:** Conceptualization, Validation, Formal analysis, Investigation, Writing – Original Draft, Writing – Review & Editing, **Anthony Piro:** Resources, **Aaron Au:** Resources, **Christopher M. Yip:** Resources, Writing – Review & Editing, **Michael B. Wheeler:** Resources **Jonathan V. Rocheleau:** Conceptualization, Writing – Original Draft, Writing – Review & Editing, Supervision

## Acknowledgments

R.R. was supported by a Natural Sciences and Engineering Research Council of Canada Postgraduate Scholarship Doctoral (NSERC PGSD). This research was supported by grants from NSERC (RGPIN-2016–371705 and RTI-2018-00846) and Canadian Institutes of Health Research (162330) to J.V.R. and stipend support to R.R. from NSERC (PGSD-535059-2019).

