## Supplemental Figures for "Islet-on-a-chip device reveals first phase glucose-stimulated respiration is substrate limited by glycolysis independent of Ca^2+^ activity"

#### Impact of SU-8 thickness on optical performance

COMSOL simulation suggests that varying the depths of the microwells does not impact the response of the oxygen sensor **(Fig. 1G-H)**. However, high numerical aperture (NA) objective lenses are commonly designed to be used with No. 1.5 glass coverslips with a refractive index of 1.515 and a thickness of 0.17 mm. Deviation from these values could induce significant spherical and chromatic aberrations. To evaluate the impact of an additional layer of the SU-8 (refractive index of 1.598) on imaging quality, we imaged fluorescent beads (MACSQuant Calibration Beads, Miltenyi Biotec) immobilized on SU-8 of varying thicknesses **(Fig. S1)**. These data show the fluorescence intensity of the beads progressively decreased as SU-8 thickness increased. Hence, a 10 μm depth was selected for the microwells to minimize the impact on imaging quality.

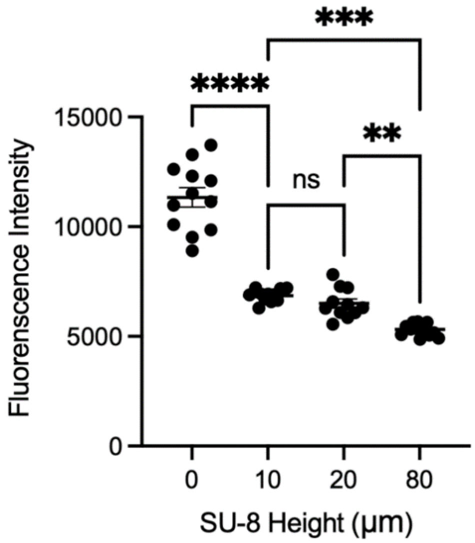


**Figure S1 Impact of SU-8 thickness on imaging quality.** A comparison of fluorescence intensities of microbeads imaged on different thicknesses of SU-8. Data in the figure are means ± SD. **** indicates $p \leq 0.0001$, *** indicates $p \leq 0.001$ and ** indicates $p \leq 0.01$ by one-way ANOVA.

#### Determining factors that affect sensitivity of $O_2$-sensors

To evaluate potential factors that could affect $O_2$ sensitivity, we modelled varying flow rates, islet proximity to the sensor and glucose-stimulated changes in $O_2$ consumption rate (**Fig. S2**). Flow rate is a key factor to consider when using PMMA channels since it is the sole means of replenishing $O_2$. Modelling various flow rates through the device showed only a moderate impact on $O_2$-sensitivity (**Fig. S2A**). These data suggest slower flow rates only marginally enhance sensor response suggesting sensing will be stable even during fluctuations in flow. Our dam-wall design can optimally trap two different sized islets; ~20 μm diameter under the dam-wall, and >80 μm diameter in the channel. Although islets sink, we were concerned that this arrangement could result in some islets not being uniformly pressed against the microwells. To evaluate the effect of proximity, we simulated islets at various heights relative to the microwells (**Fig. S2B-C**). These data show a poorer response with increasing distance between the islet and microwell, suggesting proximity is essential for sensitivity. Hence, we used a microfluidic device with channel heights that push the islet against the microwells (i.e., create a footprint). We also considered different OCR values since islets are normally studied in low and high glucose conditions. We modelled 2-

and 20-mM glucose-stimulated OCR by changing the inward flux values of the islet walls. The goal was to determine if there was a visible difference between both treatments (**Fig. S2D**). It revealed a 12.8% difference could be empirically measured. These data suggest relative changes in OCR induced by 2- and 20-mM glucose could be detected by sensor in the microwells underneath the tissue (**Fig. S2E**).

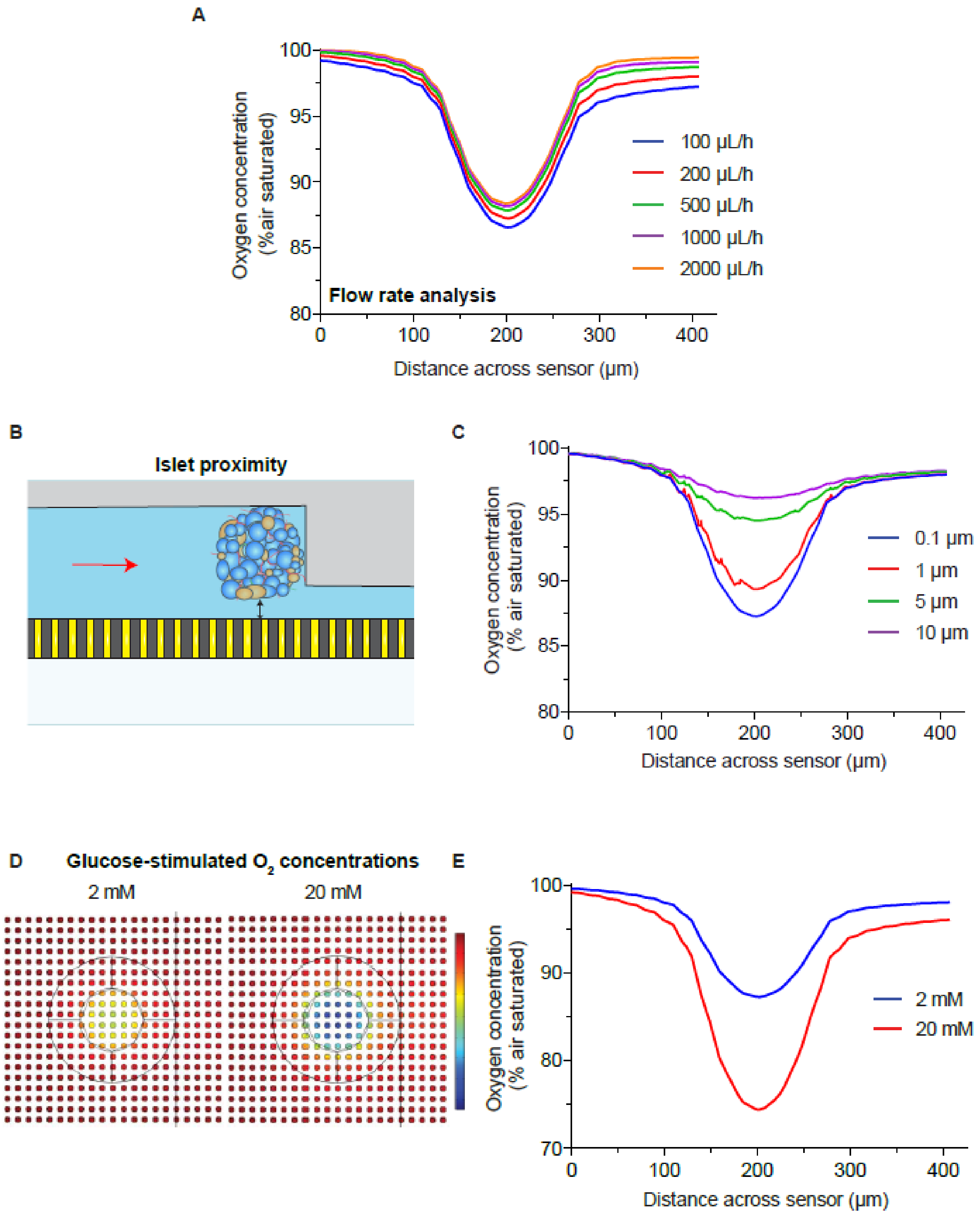


**Figure S2** Determining the impact of flow rate and islet proximity on $O_2$ sensitivity. A) Varying flow rates have minimal effect on $O_2$ levels. B) Islet proximity is measured as distance between the bottom of the islet to the top of the microwell plane. Red arrow indicates direction of fluid and $O_2$ flow. C) Simulated $O_2$ concentrations reveal proximity to enhance microwell sensitivity. D)

Glucose-stimulated $O_2$ concentrations in the wells when islets are subjected to 2- and 20-mM glucose reveal greater OCR in the center of the islet. E) Simulated $O_2$ concentrations when islets are subjected to 2- and 20-mM glucose show a large response to an increase in OCR at 20 mM glucose. Pancreatic islets were modelled as 120 μm radii cylinders with the circumferential edge filleted

### Shear stress characterization on-chip

We used dam-wall devices previously to successfully image glucose-stimulated pancreatic islet $Ca^{2+}$ activity (Rocheleau et al., 2006; Silva et al., 2013). To estimate the shear stress experienced by islets in our device, we modelled an islet trapped in a dam-wall device with a flowrate of 200 $\mu l \cdot h^{-1}$ (**Fig. S3**). These data show that most of the shear stress is localized to a small region at the bottom left and right corners of the side facing the dam-wall (**Fig. S3A**). Although the shear stress reaches values as high as 35 mPa, the histogram of relative surface area reveals that >80 % of the surface area experiences less than 5 mPa of shear stress (**Fig. S3B**). Consistent with our previous experience using these devices, dam-walled devices are relatively inert and do not significantly perturb islet metabolism and $Ca^{2+}$ activity.

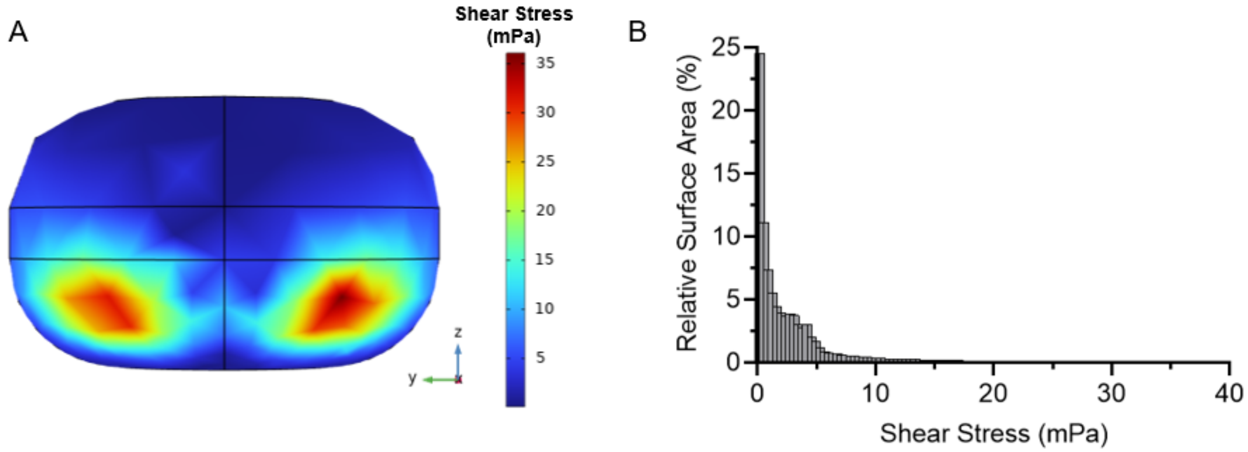


**Figure S3 Fluid flow around an islet loaded in the microfluidic device was modelled with COMSOL Multiphysics**. Islet was modelled as a 120 μm diameter cylinder with the circumferential edges filleted. A) Simulated shear stress on the surface of the modelled islet. B) A histogram of the islet surface area versus shear stress shows ~81% of the shear stress Is less than 5 mPa.

### On-chip characterization of the $O_2$ sensor

Based on linear regression, we calculated a $K_{SV}$ of 117 ± 4.8 $L \cdot mmol^{-1}$, which is consistent with literature values for this type of sensor (**Fig. S4A**) (Mao et al., 2017). The limit of detection (LOD) has previously been defined as SLM*SNR/Ksv where SLM is the uncertainty in the system's light intensity measurement, SNR is a minimum signal-to-noise ratio of 3, and Ksv is the Stern-Volmer quenching constant (Mao et al., 2017; Wang and Wolfbeis, 2014). Thus, we calculate the LOD of our sensor to be 51 $nmol \cdot L^{-1}$ based on the fitted Ksv of 117 ± 4.8 $L \cdot mmol^{-1}$ (**Fig. S4A**) and a measured SLM of ±0.2%. Our goal is to measure the dynamics of glucose-stimulated changes in islet OCR. To determine the temporal resolution of the $O_2$-sensor response, we measured the response time by switching from $ddH_2O$ to 10% sodium sulfite and vice versa (**Fig. S4B and C**). Response time, $t_{95}$, is defined as the time required to reach 95% of the luminescence maximum or minimum. These data show a $t_{95}$ of 12.4 ± 0.7 s (**Fig. S4B**), and 4.2 ± 0.2 s (**Fig. S4C**) switching from 0% to 100% air saturated $O_2$ and vice versa. These times reflect a 0.21 mM change in dissolved $O_2$ suggesting we should expect a much faster response time with the nanomolar changes induced by living islets. To measure the reproducibility of our device, we measured the response to 10% sodium sulfite in 6 different devices (**Fig. S4D**) and in a single device to repeated treatments of 10% sodium sulfite (**Fig. S4D**). These data show very good repeatability with only a 5.8 % relative standard deviation (RSD) between devices (**Fig. S4D**) and 4.2% RSD between treatments in a single device (**Fig. S4E**).

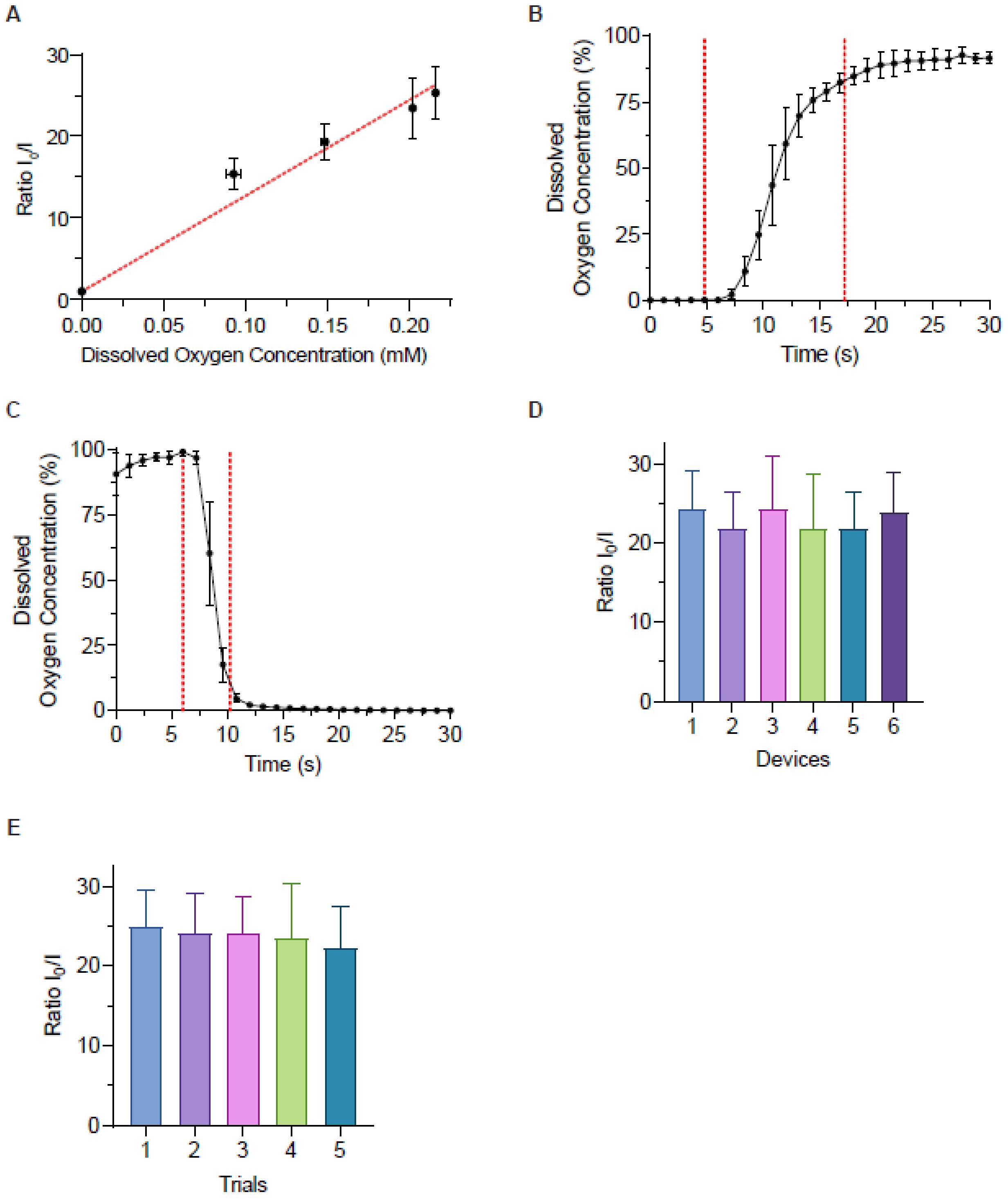


**Figure S4 Characterization of the PtTFPP/PDMS $O_2$ sensor.** A) The relationship between dissolved $O_2$ concentration and normalized phosphorescence modelled using the Stern-Volmer relationship. A representative calibration curve to calculate changes in $O_2$. N = 3 independent repeats. B) A temporal trace of switching from a 0% to 100% air saturated solution revealing a $t_{95}$ of 12.4 ± 0.69 s and an RSD of 10.6% for 6 independent repeats. Red dotted lines represent the $t_{95}$ interval. C) A temporal trace of switching from a 100% to 0% air saturated solution revealing a

$t_{95}$ of 4.24 ± 0.18 s and an RSD of 4.3% for 6 independent repeats. Red dotted lines represent the $t_{95}$ interval. D) The average $O_2$ microsensor response of 6 microfluidic devices towards 10% sodium sulfite and $ddH_2O$. E) The average $O_2$ microsensor response of one microfluidic device towards 10% sodium sulfite and $ddH_2O$ five times repeatedly. Data in the figure are means ± SD.

### Islet variability upon glucose stimulation

Due to normal islet-to-islet variability, our data analysis critically depends on measuring $Ca^{2+}$-influx and OCR in individual islets (**Fig. 3**). To show the effect of islet-to-islet variability in glucose-stimulated $Ca^{2+}$ activity, we plotted the $Ca^{2+}$ traces from 9 islets where 20 mM glucose was added at 5 min (**Fig. S5A**). These data show each islet responds to glucose with a phase 0 drop in $Ca^{2+}$ followed by a rapid phase 1 rise; however, the phase 1 rise occurred at very different timepoints (~6.5 – 9.5 min). To simulate a similar bulk $Ca^{2+}$ assay available on some commercial instruments, we also averaged the 9 traces (**Fig. S5B**). These data reveal a much more gradual response that decreases the temporal resolution of the measurement. The variability due to the different response times is particularly evident in the progressive increase in the error bars. Hence, the biphasic OCR response we measured was only possible due to our ability to measure traces of individual islets. To also show the effects of islet-to-islet variability in glucose-stimulated OCR, we simultaneously measured 4 islets in the same microfluidic device subjected to high glucose (**Fig. S5C**). These data show each islet responds to glucose by increasing the intensity of the microwells underneath the islet with some islet-to-islet variability in the magnitude of the responses suggesting the responses are independent.

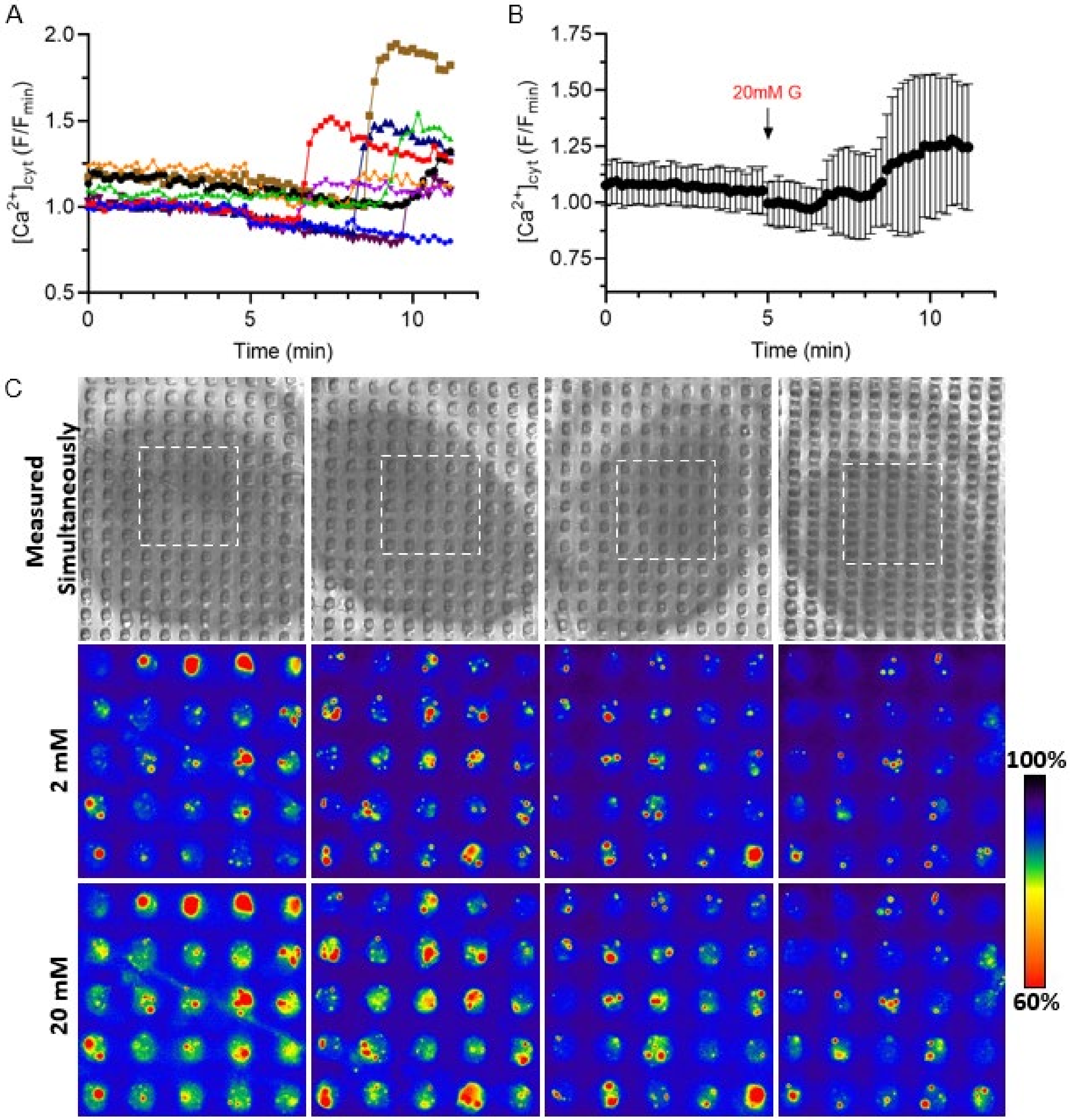


**Figure S5 Mouse pancreatic islets have variable response times and magnitudes.** A) Individual traces show variability in the timing of the responses. B) Traces in A) averaged without setting "time 0" for each curve based on the inflection points of the $Ca^{2+}$ trace. Instead, the traces are averaged based on when treatment was changed to 20 mM glucose. The arrow denotes the addition of 20 mM glucose. N = 9 islets. Data in the figure are means ± SD. C) Representative widefield white light (*top row*) and phosphorescence images of $O_2$ sensitive microwells underneath 4 islets simultaneously imaged after stimulating with 2- (*middle*) and 20-mM glucose (*bottom*) show a visible increase in phosphorescence. These simultaneous measurements illustrate variable magnitudes of respiration.

**Biphasic OCR model**

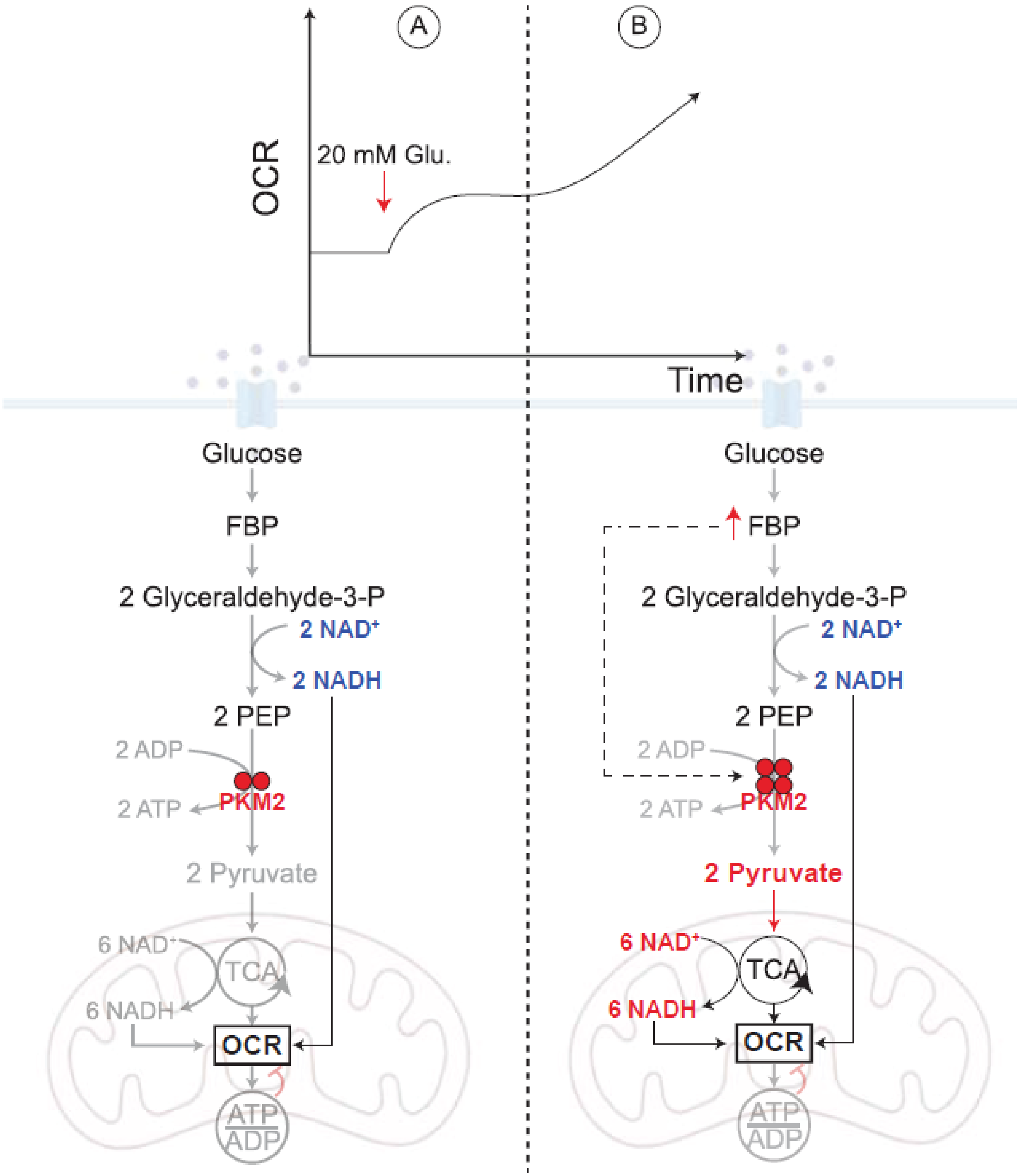


**Figure S6 A model depicting the biphasic behavior of OCR during first phase insulin secretion.** A) Upon glucose stimulation, glycolytic NADH is shuttled into the mitochondria for OxPhos, stimulating the first rise in OCR. A plateau in the OCR response follows shortly due to inactivated PKM2 (in dimer form). B) As glycolytic metabolism continues, the accumulation of upstream glycolytic metabolite, FBP, allosterically activates PKM2 (dimer to tetramer transition) causing a large pyruvate flux into the mitochondria. This pyruvate entry further stimulates TCA cycle metabolism to generate more NADH for OxPhos stimulating the second rise in OCR.
